# DeepTE: a computational method for de novo classification of transposons with convolutional neural network

**DOI:** 10.1101/2020.01.27.921874

**Authors:** Haidong Yan, Aureliano Bombarely, Song Li

**Affiliations:** School of Plant and Environmental Sciences (SPES), Virginia Tech, Blacksburg, VA 24061, USA; Graduate program in Genetics, Bioinformatics and Computational Biology (GBCB), Virginia Tech, Blacksburg, VA 24061, USA; Department of Life Sciences, University of Milan, Milan, 20122, Italy

## Abstract

**Motivation:** Transposable elements (TEs) classification is an essential step to decode their roles in genome evolution. With a large number of genomes from non-model species becoming available, accurate and efficient TE classification has emerged as a new challenge in genomic sequence analysis.

**Results:** We developed a novel tool, DeepTE, which classifies unknown TEs using convolutional neural networks. DeepTE transferred sequences into input vectors based on k-mer counts. A tree structured classification process was used where eight models were trained to classify TEs into super families and orders. DeepTE also detected domains inside TEs to correct false classification. An additional model was trained to distinguish between non-TEs and TEs in plants. Given unclassified TEs of different species, DeepTE can classify TEs into seven orders, which include 15, 24, and 16 super families in plants, metazoans, and fungi, respectively. In several benchmarking tests, DeepTE outperformed other existing tools for TE classification. In conclusion, DeepTE successfully leverages convolutional neural network for TE classification, and can be used to precisely identify and annotate TEs in newly sequenced eukaryotic genomes.

**Availability:** DeepTE is accessible at https://github.com/LiLabAtVT/DeepTE

## 1 Introduction

Transposable elements (Transposons; TEs), constitute a large portion of many known eukaryotic genomes (Makalowski, 2001; SanMiguel *et al.*, 1996), and significant roles in many biological processes (Bourque *et al.*, 2018). Accurate identification and annotation of TEs are essential to the understanding of their roles in genome evolution, genome stability and regulation of gene expression (Goerner-Potvin and Bourque, 2018; Platt *et al.*, 2016; Wicker *et al.*, 2007). With the reduced sequencing costs and novel long-read sequencing technologies, a large number of eukaryotic genomes has been sequenced in recent years. Given the diversity and abundance of TEs, annotation and classification of TEs has reemerged as a major challenge in genome annotation (Platt et al. 2016).

TEs fall into two general categories (Wicker *et al.*, 2007). Class I TEs are retrotransposons which can transpose from one to another position via the ‘copy-and-paste’ mechanism. This group contains long terminal repeat (LTR) retroelements such as Gypsy and Copia, and non-LTR retroelements such as dictyostelium intermediate repeat sequence (DIRS), penelope-like elements (PLE), long interspersed nuclear element (LINE) and short interspersed nuclear element (SINE). Class I TEs include two sub-classes. Class II TEs are DNA transposons following a ‘cut-and-paste’ mechanism, characterized by terminal inverted repeats (TIR). In Subclass 1 of Class II (Class II_sub1) TEs, nine known super families can be distinguished by the target site duplication (TSD) and TIR sequences. These TEs can be further classified into autonomous and non-autonomous elements based on their ability to move by themselves. Miniatures inverted repeat transposable element (MITE) is a special type of non-autonomous Class II_sub1 TEs with higher copy numbers and special structural feature and do not encode any transposase. Subclass 2 of Class II (Class II_sub2) TEs are DNA TEs that undergo a transposition process without double-stranded cleavage. Helitron and Maverick are two major orders in this group (Wicker *et al.*, 2007).

In the past, several computational tools have been developed to identify TEs without classifying TEs into super-families. For example, LTR TEs (LTRs) can be identified by LTR_STRUC (McCarthy and McDonald, 2003), MGEScan (Rho *et al.*, 2007), LTR_FINDER (Xu and Wang, 2007), and LTRharvest (Ellinghaus *et al.*, 2008). But these four tools cannot classify LTRs into super families such as Copia or Gypsy. MITE TEs (MITEs) can be identified by MITE-Hunter (Han and Wessler, 2010), detectMITE (Ye *et al.*, 2016), MITEFinderII (Hu *et al.*, 2018), and MITE Tracker (Crescente *et al.*, 2018). However, MITEs identified from these four tools are not classified into super families under TIR order (Wicker *et al.*, 2007). RepeatModeler is able to build and classify consensus models of putative interspersed repeats, based on sequence similarity to known repeats (Smit and Hubley, 2008). However, when we tested this method using in maize ‘B73’ genome (Schnable *et al.*, 2009), approximately 14% TEs are unclassified.

Several software have been developed for TE classification including TECLASS (Abrusán *et al.*, 2009), REPCLASS (Ranganathan, 2007), and PASTEC (Hoede *et al.*, 2014). TECLASS leverages Support Vector Machine (SVM) to classify TEs into Class I, Class II, LINE, and SINE, but only these four groups are classified. REPCLASS utilizes structural and homology characterization modules to classify TEs into more groups than TECLASS such as Class I and II, super families of LTR TEs, DNA TEs, LINE, SINE, and Helitron. REPCLASS depends on WU-BLAST which is an alignment software that is no longer maintained. An additional short-coming is that both TECLASS and REPCLASS cannot distinguish between TEs and other non-TE sequences (non-TEs). PASTEC uses Hidden Markov Model (HMM) profiles to classify TEs based on conserved functional domains of the proteins, and it identifies more super families than REPCLASS. Performance of PASTEC was shown to be better than TECLASS and REPCLASS, and it can distinguish potential host genes, rDNA, and SRR sequences from real TEs. However, the sensitivity in detecting certain groups of TEs such as TIR (64.1%), LTR (39.1%), and Class I (53.0%) groups are low, and it cannot distinguish super families under TIR, LINE, and SINE orders.

In this article, we have developed DeepTE, a deep learning method to classify TEs from non-TE sequences and to classify TEs into multiple orders and classes. Deep learning has been widely applied and has becomes one effective strategy in identifying complex patterns derived from feature-rich datasets. One particular deep learning model--Convolutional Neural Network (CNN)--have achieved outstanding performance in image classification, speech recognition, and natural language processing (Krizhevsky *et al.*, 2012; Schmidhuber, 2015). CNN model has also been successfully applied in prediction of unknown sequences profiles or motifs and functional activity discovery, without pre-defining sequence features such as prediction of sequence specificities of DNA- and RNA-binding proteins (Alipanahi *et al.*, 2015), effects of noncoding variants (Zhou and Troyanskaya, 2015), and classification of alignments of non-coding RNA sequences (Alipanahi *et al.*, 2015; Aoki and Sakakibara, 2018; Schmidhuber, 2015; Zeng *et al.*, 2016; Zhou and Troyanskaya, 2015).

Traditionally, transposons can be classified into different super families underlying distinct sequence patterns (Wicker *et al.*, 2007). In view of this, we have designed DeepTE to classify TEs into seven orders which include 15, 24, and 16 super families in plants, metazoans, and fungi respectively. This tool combines TE domain detection which improves its performance, and it outperforms PASTEC in most TE categories.

## 2 Methods

### 2.1 Transposon dataset

DeepTE was built using RepBase (Bao *et al.*, 2015) and Plant Genome and Systems Biology (PGSB) repeat database (Spannagl *et al.*, 2015). Repbase contains prototypic sequences representing repetitive DNA from 134 species. PGSB database provides access to the repeat element database from 44 species and cover 20 different genera. We combined these two datasets since TEs in these two databases are largely not overlapping. We found 74,383 TEs when combining sequences from both databases. After removing identical TEs using BLAST (Madden, 2013), there are 71,049 unique TEs in the dataset that is used for model training and testing. Species-specific (plants, metazoans, and fungi) training data were selected based on the presence of TE families in different species. For example, 63,416 TEs (out of 71,049 TEs) were used for plants (see Table S1 for a detailed breakdown of numbers of TEs in each family in plants). Because both databases only contain Helitron, the plant Class II_sub2 classifier only represent Helitron superfamily. To classify Class II_sub1 transposons into MITE and nMITE (not MITE) TEs, we manually created MITE and nMITE datasets. Class II_sub1 TEs annotated as MITE in RepBase and PGSB were labeled as ‘MITE’. The rest of Class II_sub1 TEs were annotated for PfamA domains with hmmer3 (Eddy, 2010). The TEs were defined as ‘nMITE’ where transposase (TR) domains were identified with length >= 800 bp (Wicker *et al.*, 2007) (Table S2). A total of 752 TEs were defined as ‘MITE’, and 2,696 TEs were defined as ‘nMITE’ (need something in the Table S1).

### 2.2 Transposon sequence representation

We used k-mer occurrence to construct feature matrices from the TE sequences. This system has been successfully applied to bacteria taxonomic classification of metagenomic data (Fiannaca *et al.*, 2018). The number of features was set to L = 4^k^ where k is the length of the k-mers (Figure 1). A sliding window over the input vector was used to generate the convolution operation of the first stage of the CNN network. In our analysis, we used k = 3 to 7 to test the performance of different k-mer size.

**Fig. 1.**
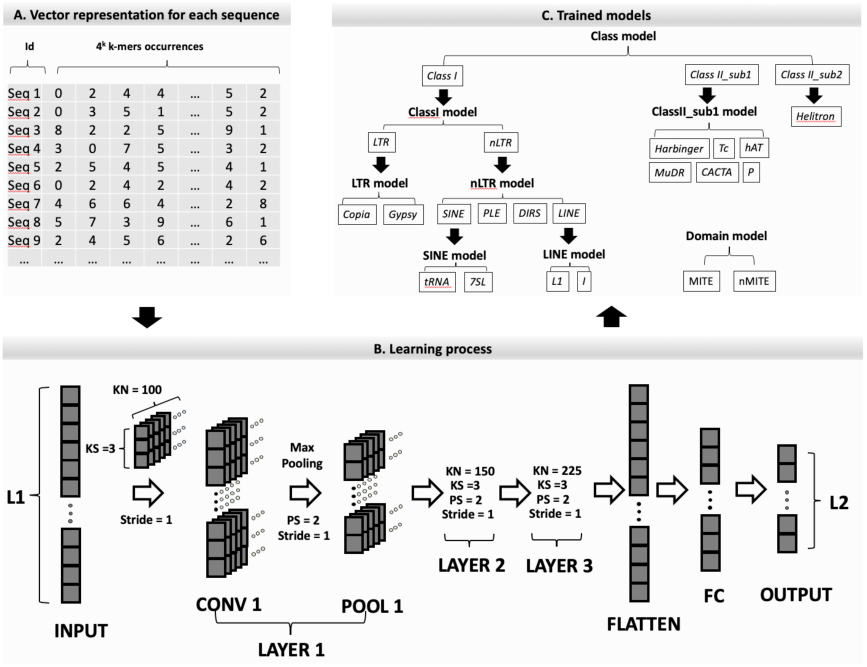
Training process of neural network. A) The training sequences were converted to a matrix of k-mers occurrences. B) The architecture of the convolutional neural network. L1 indicates the dimension of the input K-mer vector. L2 indicates the dimension of the output vector of classification number. KS represents kernel size and KN is the number of kernels. PS represents max pooling window size which is 2 and stride is equal to 1 for all three layers. FC indicates fully connected layer with 128 units. LAYER 2 and 3 had similar architectures as LAYER 1 except for number of kernels. C) Eight models (bold fonts) were trained in our classification pipeline. **Class model** was to classify TEs into Class I, Class II_sub1, and ClassII_sub2; **ClassI model** was to classify TEs into LTR and nLTR; **ClassII_sub1 model** was to classify TEs into P, Harbinger, Tc, MuDR, hAT, and CACTA; **LTR model** was to classify TEs into Copia, Gypsy; **nLTR model** was to classify TEs into SINE, LINE, PLE, and DIRS. **LINE model** was to classify TEs into L1, I; **SINE model** was to classify TEs into tRNA, 7SL; **Domain model** was to classify TEs into MITE and nMITE.

### 2.3 CNN classifier

We used a python package Keras (version 2.2.4) to implement convolutional neural network in our model (Figure 1B). Our neural network is consisted of three hidden layers with kernel sizes of three and three max-pooling layers with pool size of two (Figure 1B). The rationale of using CNN and the reasons for the choices of hyper-parameters are included in the discussion section. Hidden layers reside in-between input and output layers. For each input sequence, the k-mer frequency is calculated for different sizes of k (k = 3 to 7). In the input layer, kmers are ordered alphabetically and the orders are identical for all input sequences. Kernel is a small vector of size KS (KS = 3) where KS adjacent input kmer-frequencies are use as input for a neuron. We set the stride size as 1 which means the kernel window is moving alone the input vector with a step size of 1. Each convolution layer consisted of KN (KN = 100, 150, and 225) number of kernels. Max pooling is to transform data by taking the maximum value from the values observable in the window and max pooling size represents the height of the pooling window. Max pooling was used to summarize the output of each two adjacent kernels with a stride of 1. A dropout value of 0.5 was set after the last convolutional layer. The dropout is the probability of training a given node in a layer (0 means no output from this layer, and 1.0 means no dropout). The output of max-pooling layer is flattened as input for the next convolution layer. After the max pooling layer of the third convolution layer, a fully connected layer was set to densely connect the flattened max-pooling layer to 128 units. A dropout value of 0.5 was set after this layer. A softmax output layer was set to compute the probabilities for the classes of input sequences (Figure 1B). ReLU function was used in all three hidden layers and the fully connected layer. ReLU is an activation function that introduces non-linear properties to the network, and convert an input signal of a node to an output signal. ReLU gives an output x if x is positive and 0 otherwise (Agarap, 2018).

### 2.4 Improving classification via detecting transposon conserved domains

The eight models were organized according to TE classification system (Wicker *et al.*, 2007) (Figure S2).To improve the performance of classification, conserved domains from TE sequences were identified using PfamA domains with hummer3 (Eddy, 2010) (Table S2). Classification results from “Class” model and “ClassI” model were corrected by the following criteria: 1) If TEs were classified into Class I group with domain ‘TR’ (Transposase), the predicted ‘Class I’ was changed to ‘Class II_sub1’ in the Class model; 2) if TEs were classified into Class II_sub1 group with domain ‘RT’ (Reverse Transcriptase), this prediction was changed to ‘Class I’ in the Class model; 3) If TEs were classified into LTR but with ‘EN’ (Endonuclease) domain, this predicted group was corrected to nLTR, since ‘EN’ is exclusive in nLTR TEs (Wicker *et al.*, 2007) (Figure S2).

### 2.5 Model training and evaluation

The dataset was divided into training (95%) and testing (5%) sets for each TE family. We performed 10-fold cross validation in the training dataset, and separated the training set to sub-training (90%) and validation (10%) sets. The sub-training and validation sets were used to compare performance of different k-mer size using precision, sensitivity, and f1-score for eight models. These eight models were then used to classify TEs from the test set (Figure S2). To further evaluate the performance of DeepTE, we compared DeepTE with the latest TE classification tool called PASTEC (Hoede *et al.*, 2014). true positive (TP), true negative (TN), false positive (FP), and false negative (FN) TE number were calculated and sensitivity, specificity, accuracy, and precision were evaluated. These four scores were calculated for the test TEs using PASTEC software with default settings and compared with DeepTE. SE, SP, AC, PR, and F1 were defined as follows:

Sensitivity = TP / (TP + FN);

Specificity = TN / (FP + TN);

Accuracy = (TP + TN) / (TP + TN + FP + FN);

Precision = TP / (TP + FP);

F1 = 2 * (PR * SE / (PR + SE))

### 2.6 Classifying non-transposon and transposon sequences

To differentiate non-TE sequences from TE sequences, we collected non-transposon sequences including coding sequences (CDS) and intergenic sequences (INS). The CDS from twenty-one plant species were collected from phytozome (https://phytozome.jgi.doe.gov/index.html), and Sol Genomics Network (https://solgenomics.net/organism/Nicotiana_tabacum/genome) (Table S3). Sequences of CDS from each species were combined and CDS annotated with transposons were removed. From these CDS sequences, we randomly selected ∼800,000 sequences as our CDS dataset. These sequences were then randomly divided into ten datasets of each contained ∼80,000 sequences. For INS, sixteen plant species with known repeat annotations in the phytozome were analyzed. The intergenic sequences of each species without overlapping with transposons were collected, and sequences with length between 50 - 10,000 bp were retained. Approximately 800,000 sequences from the filtered sequences were selected, and randomly divided into ten replicated datasets with ∼80,000 for each. Each set from CDS and INS was separated into training (90%) and validation (10%) sets. TE dataset was constructed using all TE sequences (∼80,000). These TE sequences were randomly partitioned into ten equal sized subsamples, and a single subsample is retained as the validation set and the remaining nine subsamples were used as training set. This process was repeated ten times to obtain ten replicated TE datasets with each contains training and validation sets. For replicates in CDS, INS, and TE datasets, we selected one replicate from each of these datasets, and combined them to one final replicate. This process was repeated ten times and each replicate was used exactly once. Ten final replicates were generated to train model. Three scores (precision; sensitivity; f1-score) generated from keras packages were used to evaluate performances of this model.

## 3 Results

### 3.1 Performance comparison for different k-mer sizes

To identify suitable k-mer size for representing the input sequences, performances of different k-mer sizes were compared. The precision changed as the k-mer size changed in all eight trained models (Figure 2). The performance improved significantly (*p* < 0.05) with increased k-mer sizes in **Class, ClassI, ClassII_sub1, LTR**, and **nLTR** models. No significant change was found in **Domain, LINE**, and **SINE** models. Using k = 6 has similar performance with using k = 7 for most models except for **ClassII_sub1** model. In model **ClassI** and **nLTR**, using k = 5, 6 and 7 have the same performance. Using k = 3 has the worst precision across all tested k-mer sizes. Another two scores precision and f1-score showed similar results as precision (Figure S1). Taken together, using k = 7 has the best performance in all models, and it was used for further analysis. Using larger k-mer sizes requires substantial more computing time and were not tested in this analysis.

**Fig. 2.**
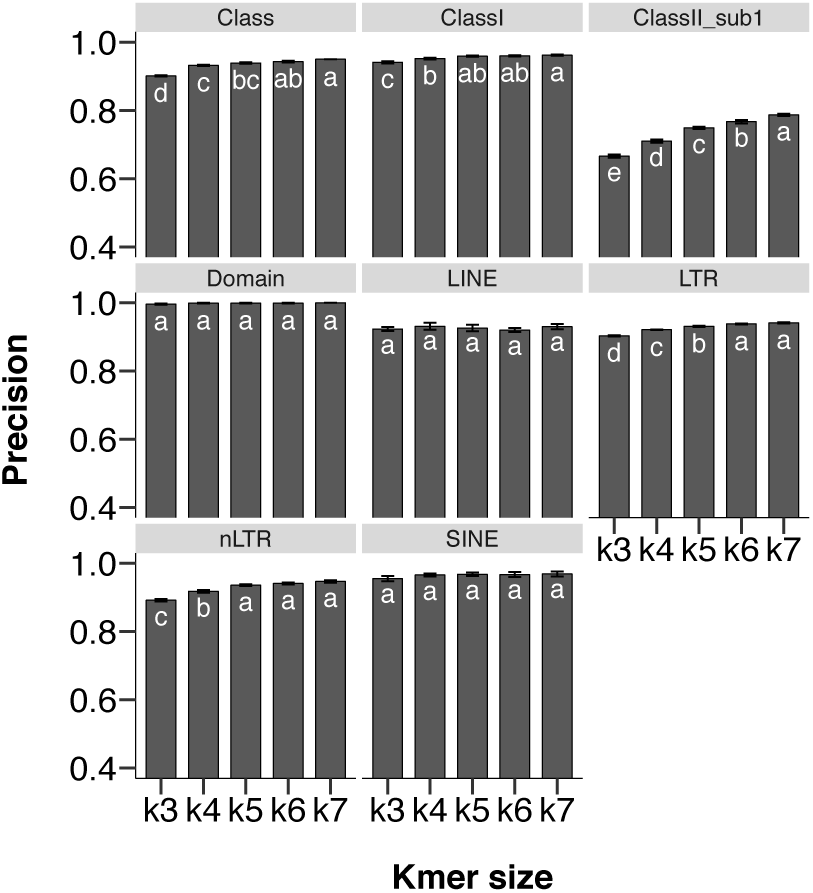
Precision validation of CNN classifier based on k-mer size. k3, k4, k5, k6, and k7 represent k-mer size from 3 to 7. Different letters insides each bar indicates significant difference with p value < 0.05.

### 3.2 Performance of trained models

To further evaluate performance from these eight models, precision, sensitivity, and f1-scores, were calculated (Table 1). In **Class** model, these three scores are higher than 0.90 for Class I and Class II_sub1 TEs. For Class II_sub2 (Helitron) TEs, although precision (0.88) is close to 0.90, its sensitivity (0.44) and f1-score (0.59) are the lowest among these three classifiers. In **ClassI** model, LTR and nLTR both have high performance, and the performance scores are all higher than 0.97 for LTR TEs. **ClassII_sub1** model could classify six DNA/TE families, with variable performance. In P super family, precision (0.50), sensitivity (0.10) and f1-score (0.18) are lower than other families. Mutator and Harbinger has higher precision than sensitivity and f1-score. For TcMar and hAT, their sensitivities (> 0.80) are higher than precision (< 0.80) and f1-score (< 0.80). CACTA has relative consistent scores with average of 0.70. For **LTR, nLTR, LINE** models, all super families have similar performance scores, and most scores are above 0.90. Particularly, **Domain** model displayed the best performance with MITEs and nMITEs achieved to maximal value for these three scores (Table 1).

**Table 1.**
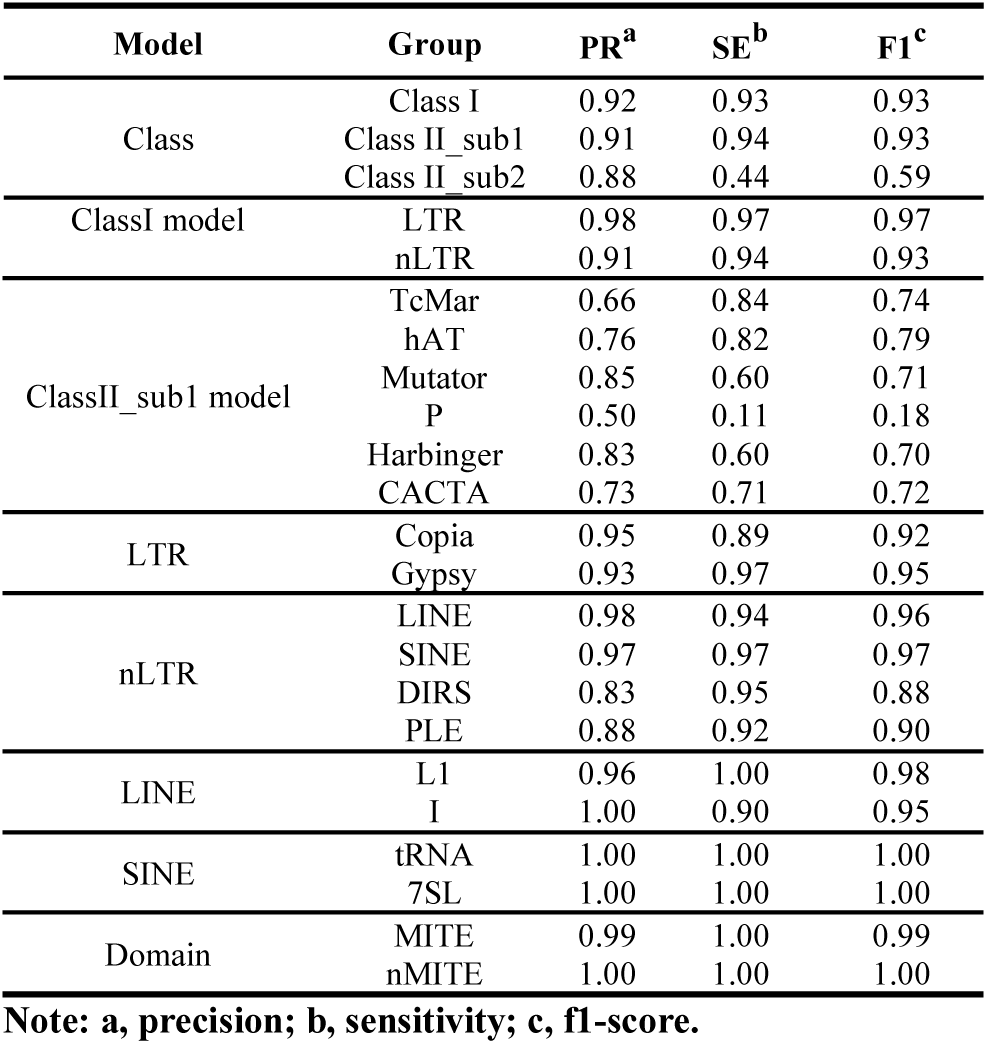
Performance of eight models in plants.

In summary, we can roughly divide the models into four categories: high, good, moderate and poor performance. High performance models have all three scores higher than or equal to 0.9. Moderate high models have at least one score higher than or equal to 0.9 and all scores are higher than 0.7. Moderate models have no more than two scores that are lower than 0.7. Poor models have all three scores below 0.7. Among all TE classification results, we have 50 percent high performance models (4/8), 25 percent good models (2/8), one moderate model and one poor models.

### 3.3 Improving classification by discovering conserved domains of transposon sequences

These eight models were wrapped together to classify unknown TEs and the classification performance were evaluated with four scores including accuracy, precision, sensitivity, and specificity (Figure S2; Figure S3). DeepTE showed high accuracy over 0.90 for all TE groups, and more than half of them (61%; 14/23) have almost perfect accuracy over 0.98 (Figure S3A). In Class I, all nLTR groups showed higher accuracy than all LTR groups. Among all TE groups, 70% (16/23) had SE higher than 0.80, and most Class I groups (92%; 12/13) had sensitivity over 0.80. ClassII_DNA_P showed the least sensitivity (0), probably due to its low number in the dataset (Table S1). ClassI_nLTR_SINE_7SL, MITEs and nMITEs had the highest sensitivity (Figure S3B). For precision, over half of TE groups (57%; 13/23) were over 0.70 (Figure S3C), while for specificity, nearly all groups were over 0.90 (Figure S3D). Taken together, our method showed higher sensitivity for Class I groups than Class II_sub1 and Class II_sub2 groups. For accuracy, precision, and specificity, our method showed similar performance in all TE groups. Remarkably, for Class II_sub1 MITE and nMITE groups, our method generated sensitivity, accuracy, precision, and specificity close to 100%. Our method showed lower sensitivity for class II_sub2 than the other TE groups, but it performed better in accuracy, precision, and specificity for Class II_sub2.

We can also roughly divide the 23 TE groups into four categories as defined in the performance of trained models: high, good, moderate and poor. In total, we have 17.4 percent high performance models (4/23), 34.8 percent good models (8/23), and 47.8 percent moderate models (11/23).

TE families were defined with high DNA sequence similarity for their typical internal domain or coding region (Wicker *et al.*, 2007). To improve performance of TE classification in DeepTE, conserved domains of TEs were detected to correct false classification (Figure S2; Table S2). Several groups show slightly improved accuracy, sensitivity, precision, and specificity; however, these improvements are not statistically significant (Figure S4; Table S4).

In general, these eight models achieved good performance, particular for accuracy and specificity, and it had relative better sensitivity score in Class I, than in Class II_sub1 and II_sub2 families. The classification performance was slightly increased after correction of false prediction *via* detecting domains within TE sequences.

### 3.4 Performance comparison between DeepTE and PASTEC

DeepTE was compared with a published TE classifier called PASTEC. There are 11 TE groups that can be classified by both DeepTE and PASTEC, and the model performances were used in our comparison, including eight Class I groups, three Class II_sub1 groups, and Class II_sub2 (Figure 3).

**Fig. 3.**
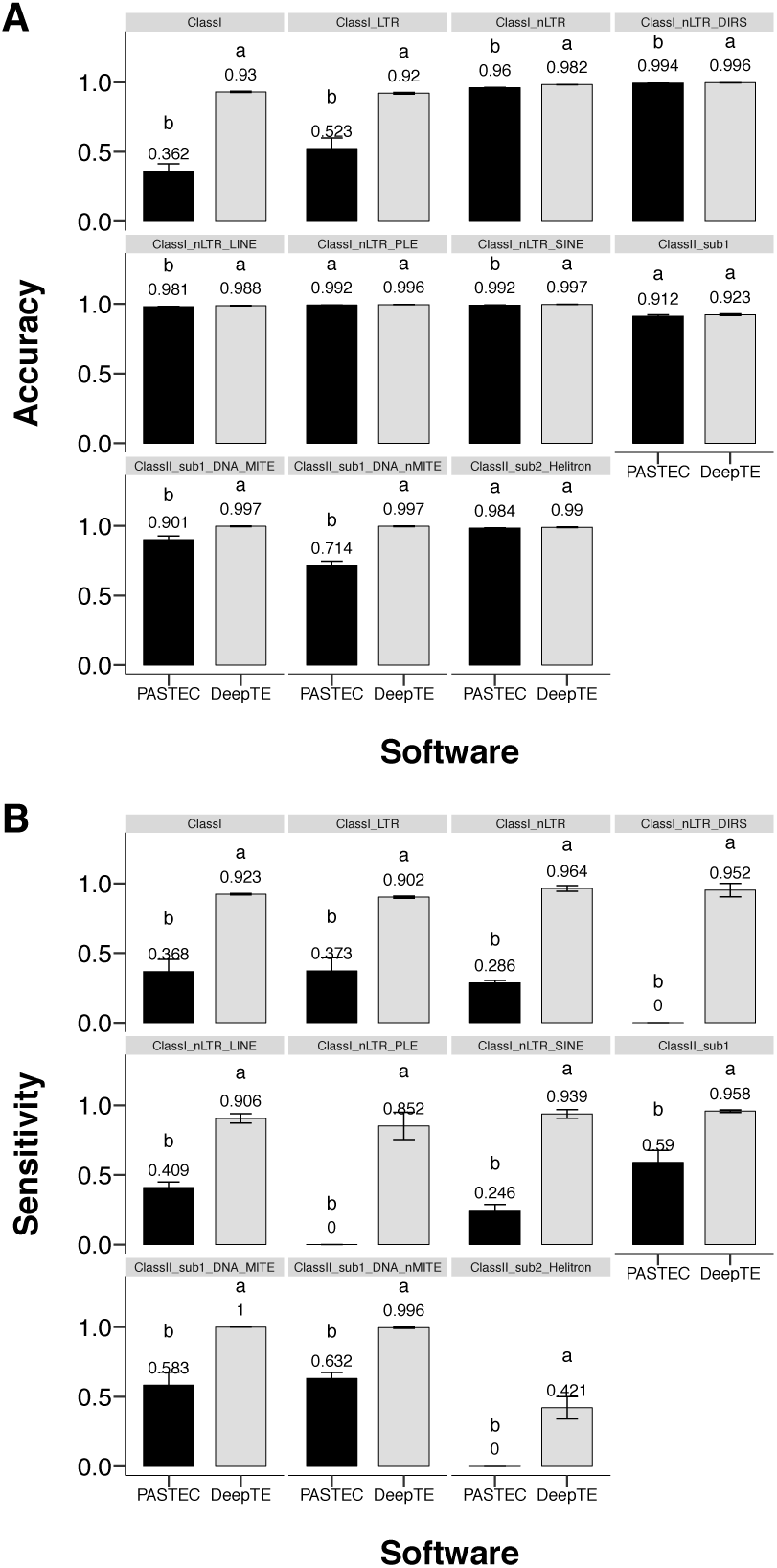
Performance comparison between DeepTE and PASTEC. Different letters above each bar indicates significant difference with p value < 0.05.

In comparison of accuracy, PASTEC showed lower performance as compared to DeepTE for most groups, except for Class II_sub1, Class II_sub2, and PLE where PASTEC showed no significant difference than DeepTE (*p* value > 0.05). DeepTE had more than two folds accuracy score than PASTEC in Class I, and more than 1.5 folds accuracy score in LTR (Figure 3A). PASTEC showed lower sensitivity in all TE groups, of which the highest sensitivity was smaller than 0.65, and no TE was detected in DIRS, PLE, and Class II_sub2 groups. The sensitivity performance of all the groups were better in DeepTE than in PASTEC. In DeepTE, most groups (10/11) had sensitivity higher than 0.85, except for Class II_sub2 (sensitivity = 0.421) (Figure 3B). PASTEC had higher precision in four TE groups, but the four groups were lower than DeepTE, and three groups showed no significant differnece. The precision in DIRS, PLE, and Class II_sub2 was 0 in PASTEC (Figure S5A). For specificity, seven groups in PASTEC displayed better performance than DeepTE, except for Class I in DeepTE with more than three folds specificity than PASTEC. No significant difference was found in MITE, nMIT, and Class II_sub2 (Figure S5B). Taken together, PASTEC detected low number of false positives, resulting in better precision performance than DeepTE in LTR, nLTR, LINE, SINE, and Class II_sub1 groups. But it failed to detect plenty of true positive TEs, even it did not find any true positives in DIRS, PLE, and Class II_sub2. Taken together, DeepTE outperforms PASTEC.

To further compare DeepTE and PASTEC, receiver operating characteristic (ROC) curves for DeepTE were generated. Because PASTEC cannot generate ROC curves, we mark the performance of PASTEC on the DeepTE ROC curve as individual data points (Figure S6). In DeepTE, except for ClassII_sub1, all models have area under curve close to 1, suggesting good performance for these models. In PASTEC, all groups showed false positive rate close to 0 except for Class I, but their true positive rates were all below 0.65, and Class II_sub2, DIRS, and PLE groups showed 0 true and false positive rates.

We also compared the speed for DeepTE and PASTEC. The computing time for DeepTE and PASTEC were tested on a Linux workstation with Inter(R) Xeon(R) Gold 5115 CUP (2.40 GHz), eight cores and 40 GB memory. There is one graphic card installed in this Linux workstation: NVIDIA Corporation GP102GL [Tesla P40] (1.30 GHz base and 1.53 GHz as boosters) with 24 GB memory. For DeepTE, the time used to classify 1,084 TEs was 71 ± 3 seconds, and for PASTEC, the time used to classify these TEs was 1,305 ± 7 seconds. These results suggest that DeepTE is 18.3 times fasters than PASTEC when classifying TEs.

### 3.5 Classification of non-TE and TE sequences in plants

As an additional function for DeepTE, we have trained a model to classify CDS, INS, and TE datasets in plants to differentiate non-TE and TE sequences. Three scores (precision, sensitivity, and f1-score) were used to evaluate model performance. The f1-score in CDS was significantly higher than TE and INS groups, and no difference was found between TE and INS groups. For precision, score in INS was lower than scores in TE and CDS groups. CDS showed the highest score in sensitivity, while the score in TE was the lowest (Figure 4). In total, CDS group achieved the best performance comparing with TE and INS groups.

**Fig. 4.**
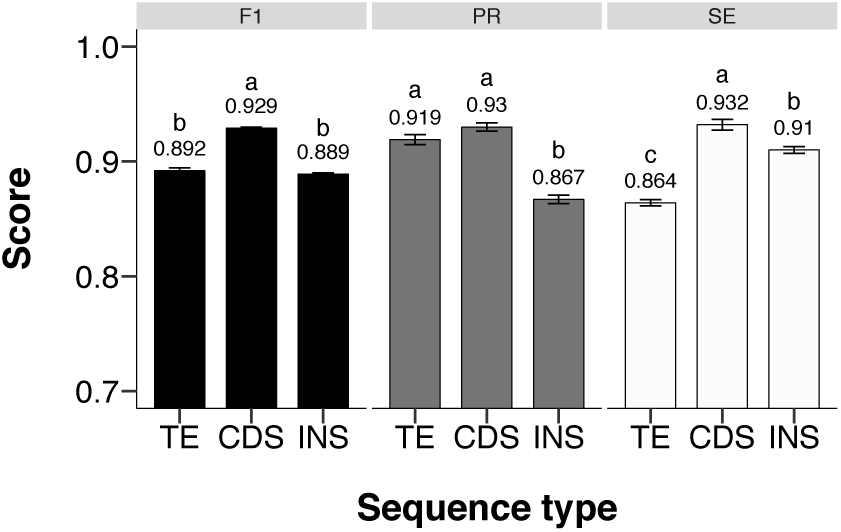
Comparison of performance for distinguishing non-TE and TE sequences. CDS: coding sequences. INS: intergenic sequences. F1: f1-score. PR: precision. SE means sensitivity. Different letters above each bar indicates significant difference with p value < 0.05.

### 3.6 TE classification in metazoans and fungi

Metazoans and fungi have TE super families that are not detected in plants, we developed another two training datasets containing 71,049 and 63,449 TEs in metazoans (Table S5) and fungi (Table S6), respectively. Seven models were trained to classify TEs into 24 and 16 super families in metazoans and fungi, respectively, based on same model structures as plants (Table S7-S8). Consistent with the plant results, metazoans and fungi both showed the worse performance in ClassII_sub1 than in other models, and most groups (66%; 21/32) in metazoans and (67%; 16/24) in fungi had evaluation scores over 0.7 of precision, sensitivity, and f1-score (Table S7-S8).

## 4 Discussion

Many studies applied deep learning models to predict unknown genomic sequences directly using sequence information, instead of pre-defined features (Eraslan *et al.*, 2019; Park and Kellis, 2015). CNN is an essential model of deep learning, and suitable for identifying sequence profiles, due to its excellent feature extraction capability on high-dimensional data (Kelley *et al.*, 2016; Zeng *et al.*, 2016). The input vector of CNN is primarily based on sequence-derived features, such as the frequency of k-mer occurrence applied in this study and one-hot vector strategy (Aoki and Sakakibara, 2018; Fiannaca *et al.*, 2015; Ghandi *et al.*, 2014; Lee *et al.*, 2011; Nguyen *et al.*, 2016). One apparent advantage of the one-hot vector is to reserve specific position information of each individual nucleotide in sequences. In the case of TE classification, because TEs are different in their sizes, one-hot vector is not directly applicable. Alternatively, the k-mer approach can be directly applied to sequences with different sizes and the location of the k-mer inside a sequence does not need to be fixed (Eraslan *et al.*, 2019). This method fits TE structure well, since TEs are classified different families commonly based on motif categories or flanking sequence patterns, which can be captured by k-mers (Wicker *et al.*, 2007).

We tested different hyper-parameters including the number of network layers, kernel sizes, kernel numbers, and k-mer sizes. However, we found these hyper-parameters have small impact on the model performance except for the k-mer sizes. We have compared performances of different k-mer sizes, and found that using k = 7 performed best. We also tested eight and higher k-mer sizes, but the time it takes to train the model is prohibitive, as for the k-mer number exponentially increases with k-mer length. In ClassII_sub1 model, we did observe a trend where k = 7 showed better performance than k = 6. (Figure 2). However, precision of LINE, SINE, and Domain models did not increase as k-mer size increased. Class, ClassI, LTR, and nLTR models did not show consistent improvement as k increases. This observation suggests the precision may not be sensitive to variations of k-mer size. For example, in the LINE model, L1 and I super families could be clarified since the I superfamily has an additional RH domain compared with the L1 superfamily. In contrast, domains of all Class II_sub1 super families are transposase, resulting in significant different performances among all five k-mer sizes (Figure 2).

Currently, more than 27 and 17 TE super families are identified, but our study can only classify TEs into 24 and 16 super families in metazoans and fungi (Wicker *et al.*, 2007). The missing families were not considered because there is only small number of TEs in the databases. In plants, there are only 15 known super families and DeepTE can classify all these TE super families. However, the collected super families have varied number of TEs, potentially reduces the classification accuracy (Barandela *et al.*, 2004). For example, the number of TE/Gypsy (30,368) is 150 times more than the number of TE/P in plants (Table S1). To handle this imbalanced issue, we leveraged a tree structural classification process (Figure 1) by setting eight models that could classify the super families under each order. Among these models, performance of ClassII_sub1 was lower to the others (Table 1). This could be accounted for most Class II_sub1 super families are distinguished by sequence dissimilarities of the terminal inverted repeats and TSD size, rather than motifs easily defined in TE body region (Wicker *et al.*, 2007), possibly raising difficulties to identify the sequence differences.

## 5 Conclusion

With the growing availability of reference genome from non-model species, TE annotation becomes a new challenge. In this study, we developed a deep learning tool called DeepTE to classify unknown TEs based on convolutional neural networks, and it outperforms current PASTEC tool. DeepTE contained eight models for different classification purposes, and also wrapped a function to correct false classification based on domain structure. This tool classified TEs into 15-24 super families according to the sequences from Plants, Metazoans, and Fungi. For unknown sequences, DeepTE could distinguish non-TEs and TEs in plant species. A manual is provided for users to apply DeepTE in identification and annotation of TEs in a different genome (Supplementary Note). This tool provides a new method for using deep learning on TE identification or annotation.

## Acknowledgements and Funding

This work is supported by USDA Hatch program and translational plant sciences program at Virginia Tech.

